# Efficient permuted intron-exon RNA circularization system based on the LtrB group II Intron from *Lactococcus lactis*

**DOI:** 10.64898/2026.10.20.760490

**Authors:** Kathrina D. Castillo, Chotiwat Seephetdee, Wataru Horikawa, Daniel L. Kiss

## Abstract

**Key findings:** Circular RNAs (circRNAs) have emerged as promising platforms for stable RNA expression and therapeutic applications, but methods for their generation remain limited. Here, we engineered the *Lactococcus lactis* LtrB group II intron into a permuted intron-exon (PIE) configuration to enable efficient circRNA formation (circLtrB). Among the tested split designs, LtrB domain IV cleavage yielded the highest circularization efficiency. Although LtrB encodes LtrA maturase required for efficient *in vivo* splicing, *in vitro* application of LtrA to the PIE constructs resulted in lower circularization efficiency than self-splicing conditions. We further examined circularization under distinct salt conditions and showed that NH_4_Cl- and KCl-based splicing buffers independently support high circularization efficiencies (>80%), while producing distinct splicing intermediates, indicating ion-specific mechanistic effects on circularization. Extending outer homology arms did not enhance circularization efficiency, demonstrating that efficient circRNA formation in the circLtrB platform is independent of extended exon pairing interactions. Using this system, we generated circRNAs across a range of coding sequence lengths, highlighting its payload flexibility. To optimize translation, we evaluated multiple internal ribosome entry sites (IRES), including Aichivirus, Crohivirus, CVB3, improved EMCV, HRV, and Salivirus, and found that the CVB3 IRES drove the highest translation efficiency. Linear NLuc showed faster decay kinetics, while circLtrB-NLuc exhibited more sustained expression over time. Transcriptome-wide RNA sequencing revealed comparable innate immune responses between circular and linear RNAs. Together, these results establish circLtrB as a versatile and efficient platform for circRNA production, providing mechanistic insight into group II intron-mediated circularization and expanding the toolkit for functional circRNA engineering.

## INTRODUCTION

RNA-based therapies are a rapidly evolving and transformative class of medicines, complementing traditional small molecules, biologics, and cell-based treatments in addressing a broad range of diseases. While early RNA therapeutics primarily focused on messenger RNAs (mRNAs), small interfering RNAs (siRNAs), and anti-sense oligonucleotides (ASOs), circular RNAs (circRNAs) have recently gained attention as a distinct platform for therapeutic RNA engineering (Wesselhoeft et al. 2018; Kristensen et al. 2019; Tong et al. 2025a). circRNAs were first discovered in the 1970s in the form of plant viroids, and later identified in eukaryotes in the early 1990s (Sanger et al. 1976; Arnberg et al. 1980; Nigro et al. 1991). Naturally occurring circRNAs are generated through backsplicing, which yields covalently closed RNA molecules that resist exonuclease-mediated degradation due to the absence of exposed 5′ and 3′ ends (Wilusz and Sharp 2013; Enuka et al. 2016; Zhang et al. 2016). Naturally-occurring circRNAs are now identified through high-throughput sequencing and advanced bioinformatics as abundant, predominantly non-coding transcripts in diverse organisms (Salzman et al. 2012; Memczak et al. 2013). circRNAs have been reported to function as microRNA or protein sponges, scaffolds for protein complexes, regulators of mRNA translation and stability, and biomarkers for various diseases (Hansen et al. 2013; Li et al. 2015; Du et al. 2016; Abdelmohsen et al. 2017; Chen et al. 2021). Although the majority of endogenous circRNAs lack protein-coding potential, engineered coding circRNAs typically incorporate internal ribosome entry sites (IRESs) or other cap-independent translation elements to enable efficient protein expression (Wesselhoeft et al. 2018; Wesselhoeft et al. 2019; Prats et al. 2020).

In recent years, both coding and non-coding circRNAs have developed into promising clinical platforms, with possible applications including CAR T cell therapy, gene editing, gene therapy, and vaccines (O’Leary et al. 2025). *In vitro* circRNA synthesis can be achieved by chemical ligation, enzymatic ligation, or ribozyme-mediated circularization (Wasinska-Kalwa et al. 2025). Most ribozyme-based approaches employ group I introns, primarily in two designs: the permuted intron exon (PIE) system where intron halves flank the exons, and the self-targeting and splicing (STS) system where the intron precedes the circularized exon. Both proceed via a two-step phosphotransferase reaction initiated by GTP attachment at the 5′ exon-intron junction, followed by ligation of the released 5′ exon to the 3′ exon-intron junction. Both methods generate circRNAs that retain ligated exon sequences (Lee et al. 2024). The *Anabaena* pre-tRNA intron, the T4 bacteriophage *td* intron, and the *Tetrahymena* group I intron are all group I introns frequently used for PIE circularization (Ford and Ares 1994; Wesselhoeft et al. 2018; Cui et al. 2023; Lee et al. 2023). Engineered group I ribozymes can achieve circularization efficiencies as high as 90%, although linear RNA contaminants must be removed. For example, excised intronic sequences and a sliced open formerly circRNA (which can be up to 30% of the RNA) species in current designs may elicit immunogenic responses (Ford and Ares 1994; Wesselhoeft et al. 2018; Lee et al. 2023).

In contrast, group II introns self-splice through two transesterification steps mechanistically related to the spliceosome where the branch-point adenosine attacks the 5′ splice site to form a lariat intermediate, followed by exon ligation at the 3′ splice site and release of the intron lariat (Peebles et al. 1986; van der Veen et al. 1986). Despite offering higher precision in exon ligation compared to group I introns, group II introns remain less explored for circRNA production (Murray et al. 2001; Lambowitz and Zimmerly 2011). To date, the only explicitly reported group II intron used in a PIE-based design is that derived from *Clostridium tetani* (Wei 2025). Early work showed that modular rearrangement of the *C. tetani* intron structure could generate circRNAs with minimal or no retained sequences, achieving more than 50% efficiency across most genes and up to 60-90% with optimized junction designs (Wei 2025). Building on the *C. tetani* PIE system, a more recent study incorporated untranslated regions (UTRs) from *Flaviviridae*, which boosted circularization efficiency from ∼50% to ∼90%, comparable to *Anabaena* group I introns.

The *Lactococcus lactis* LtrB group II intron has been extensively studied as a model for RNA splicing. The LtrB intron falls within the group II category of introns, comprising six structural parts labelled D1 through D6 (Fig. 1). D1, the largest domain, hosts multiple brief exon binding sites (EBS) that interact with intron binding sites (IBS) at splicing junctions. Splice-site specificity is primarily governed by IBS1/EBS1 and IBS3/EBS3, while IBS2/EBS2 is often dispensable (Lambowitz and Zimmerly 2011). Domains 2 and 3 (D2 and D3) play crucial roles in constructing the active intron structure and initiating the splicing process. Domain 4 (D4) houses the LtrA maturase open reading frame (ORF), Domain 5 (D5) contains the active site for self-splicing, and Domain 6 (D6) includes the bulged adenosine functioning as the branching site (Fig. 1) (D’Souza and Zhong ; Matsuura et al. 2001). The LtrB intron is capable of self-splicing *in vitro* under high monovalent salts (KCl or NH_4_Cl) and divalent magnesium ions where the RNA folds into its catalytically active conformation without protein assistance. *In vivo*, efficient and accurate splicing typically requires its intron-encoded protein (IEP) maturase, LtrA. LtrA binds the intron RNA, stabilizes its tertiary structure, and promotes proper alignment of splice sites, greatly enhancing splicing efficiency under physiological conditions.(Matsuura et al. 1997; Wank et al. 1999; Belhocine et al. 2007). More recently, due to its reverse transcriptase (RT) function, the LtrB intron has been re-engineered as a precise gene-targeting vector conceptually like the CRISPR-Cas system (Enyeart et al. 2014; Velázquez et al. 2022). These applications highlight the versatility of LtrB as an RNA-based tool.

**Figure 1.**
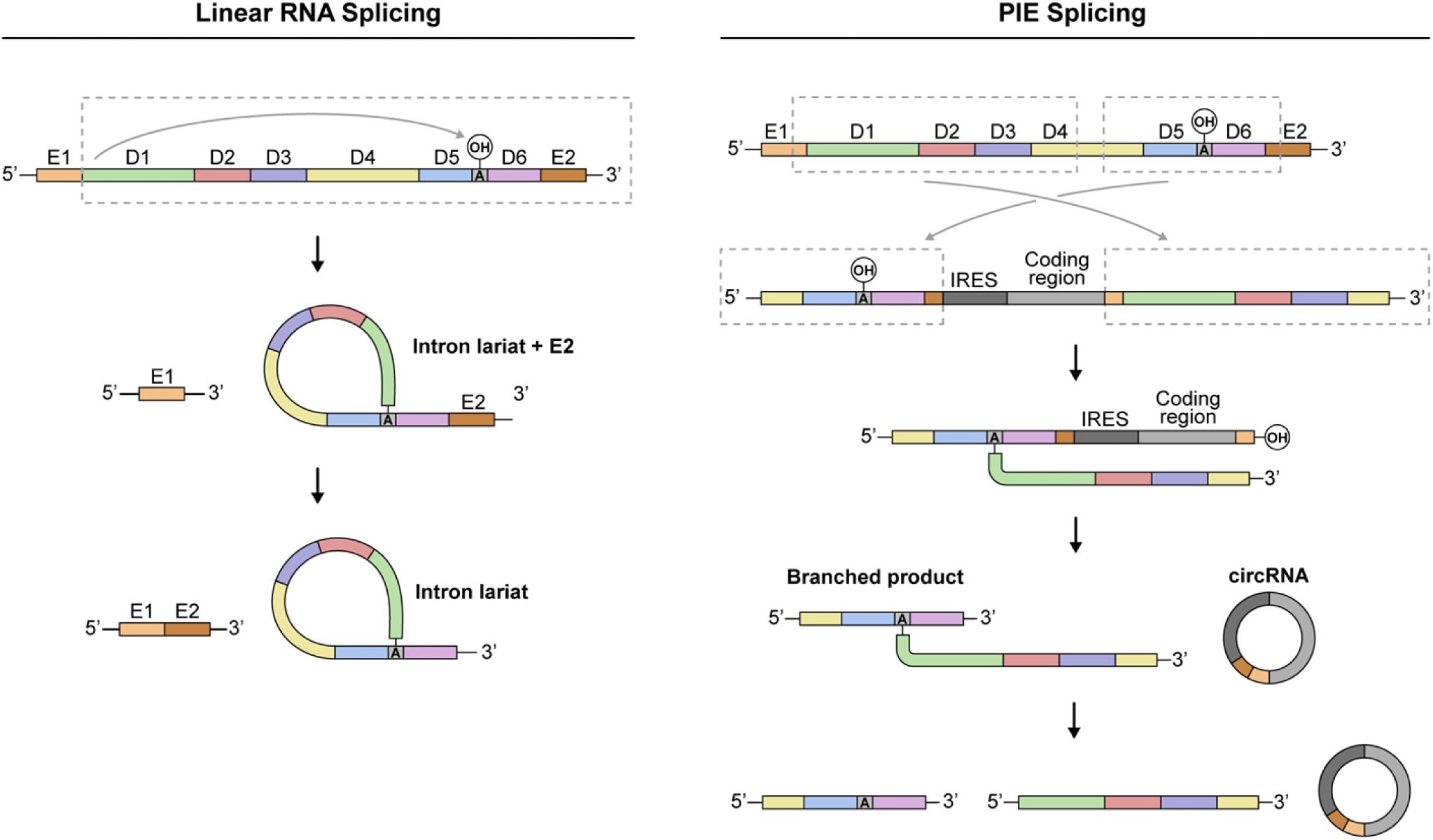
Schematic diagrams showing the typical group II linear LtrB intron and PIE construct splicing mechanisms. LtrA maturase facilitates linear LtrB splicing *in vivo* but is not required for efficient *in vitro* splicing and PIE-mediated circularization.

In this study, we explored a PIE-based strategy with the LtrB group II intron to generate circRNAs *in vitro*, using both self-splicing and LtrA maturase-assisted approaches. As this is the first use of the LtrB intron as a PIE platform, we systematically evaluated split intron configurations and reaction buffer conditions to identify parameters that maximize circularization efficiency. Among the designs tested, splitting within D4 consistently produced the most efficient circularization and was therefore selected for subsequent experiments. Using this optimized design, we show that LtrA maturase-assisted splicing yields approximately ∼25% circRNA, whereas self-splicing conditions achieve ∼90% circularization efficiency. We further demonstrate that the platform supports efficient circularization across RNA lengths, spanning ∼500 nt to ∼2,500 nt, and that circularization can still be achieved for longer transcripts (∼6000 nt), albeit with reduced efficiency. Consistent with canonical linear LtrB intron splicing, PIE-mediated splicing also required elevated concentrations of monovalent salts NH_4_Cl or KCl, underscoring the importance of ionic conditions in supporting intron folding and catalysis. CircLtrB-NLuc and linear NLuc expression were examined in HEK293 cells and monitoring over 12 days showed that CircNLuc exhibited slower decay and more sustained expression than linear NLuc. Collectively, these findings establish the split LtrB intron as a versatile and tunable alternative platform for the *in vitro* synthesis of both coding and non-coding circRNAs.

## Results and Discussion

### Engineering the *LtrB* group II intron into a PIE configuration enables circular RNA formation

Multiple strategies for circRNA generation have been described but existing platforms often exhibit cargo-dependent performance, require extensive optimization, or produce heterogeneous reaction by-products (Zhang et al. 2025). These limitations highlight the need for additional robust circRNA platforms capable of supporting efficient and scalable circularization across diverse RNA sequences and lengths (Lee et al. 2024). Group II introns, with their intrinsic autocatalytic splicing activity and capacity to accommodate diverse RNA inserts, represent a promising but underexplored system for circRNA generation.

We used the group II intron LtrB’s self-catalyzed splicing reaction to generate circular RNAs *in vitro*. The LtrB intron is comprised of six conserved domains (D1-D6), with D1 mediating splice-site recognition, D2-D3 supporting intron folding, D4 encoding the LtrA maturase, D5 forming the catalytic core, and D6 containing the branch-point adenosine required for splicing (de Lencastre and Pyle 2008) (Fig. 1). Inspection of the full-length LtrB sequence identified an early T7 polymerase termination site on D1, leaving a shorter IVT RNA product (∼400 nt) in addition to the full length LtrB RNA (Supplemental Fig. 1). A paired two-base mutation eliminated the pausing and maintained the loop structure enabling efficient production of a full length LtrB *in vitro-*transcribed RNA (Supplemental Table 1, Supplemental Fig. 1).

Using the modified LtrB intron sequence and leveraging prior studies on LtrB splicing requirements, we developed LtrB permuted intron-exon (PIE) versions. These sequences consisted of shortened exons flanked by two divided introns, yielding PIE constructs P1, P2, and P3 (Fig. 2A). Initially, the design incorporated two short 6-nt sequences, IBS3 and IBS1, within the exons at each splicing junction. These sequences were included to enable long-range interactions with the EBS3 and EBS1 of the intron. To help re-form the splicing apparatus, 20 nt homology sequences were added to the 5’ and 3’ exons (Wesselhoeft et al. 2018). Furthermore, these test constructs carried the Coxsackievirus B3 (CVB3) IRES and the nanoluciferase (NanoLuc) ORF as cargo to validate the PIE design (CircLtrB-NLuc), and to facilitate the convenient assessment of translation from the resulting circRNAs. To ensure efficient IRES-mediated translation, a 34 nt permissive spacer and a 30 nt polyA-C region have been added before the CVB3 IRES (Fig. 2A, Supplemental Table 1) (Wesselhoeft et al. 2018). In addition to the PIE versions, we also designed a linear LtrB intron with a truncated D4 (Lin1) as a splicing control (Fig. 2A, Supplemental Table 1).

**Figure 2.**
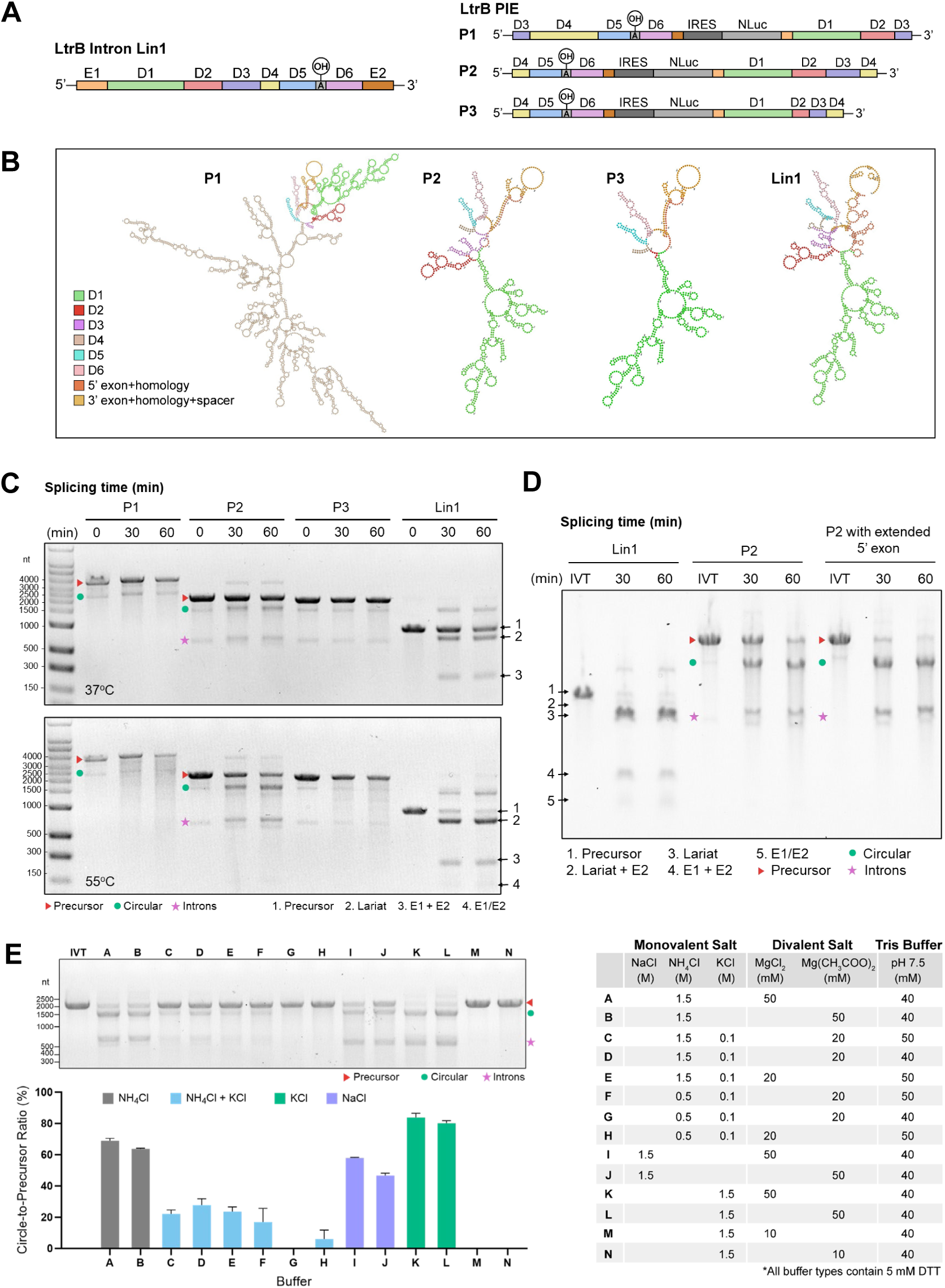
Permuted Intron-Exon (PIE) splicing of group II LtrB intron facilitates efficient RNA circularization. *(A)* LtrB construct designs showing intron split positions used to generate PIE variants; a truncated linear intron (Lin1) was included as a splicing control. *(B)* RNAfold-predicted secondary structures of constructs in *(A)* (NLuc sequence omitted). *(C)* Self-splicing of linear and PIE RNAs in standard self-splicing buffer SSB (0.5 M NH_4_Cl, 37°C) or modified SSB (1.5 M NH_4_Cl, 55°C), each containing 50 mM MgCl_2_, 40 mM Tris-Cl pH 7.5, and 5 mM DTT. *(D)* Splicing of Lin1, PIE P2 (CircLtrB-NLuc), and P2 containing a 6-nt 5′ exon extension in modified SSB. *(E)* Buffer optimization for CircLtrB-NLuc self-splicing at 55°C for 30 min (n = 3). Circularization percentages were calculated from purified circularization reactions (IVTC) and expressed relative to the remaining linear precursor.

To evaluate splicing efficiency, we initially used the original self-splicing buffer (SSB, 0.5 M NH4Cl) and incubation at 37^°^ C, as described previously (Matsuura et al. 1997). Under these conditions, we observed minimal circularization for the PIE variants, while Lin1 exhibited approximately 50% splicing after 60 min (Fig. 2C). In more recent studies, increasing the NH_4_Cl concentration to 1.5 M (Wank et al. 1999) and raising the temperature to 55°C consistent with temperatures used in published PIE circularization protocols (Wesselhoeft et al. 2019), significantly enhanced LtrB splicing efficiency. Applying these modified conditions to our system led to approximately 50% circularization efficiency for P2 and > 50% splicing efficiency for Lin1 after 60 min (Fig. 2C). This is evidenced by increased intensity of the circular RNA band, a corresponding reduction in the linear precursor, and the appearance of bands consistent with released intron. In contrast, variants P1 and P3 showed no notable improvement and exhibited ‘smeared’ bands, consistent with partial degradation (Fig. 2C). The better performance of P2 may reflect similarities between the predicted secondary structures of P2 and the linear Lin1 construct (Fig. 2B), suggesting that proper RNA folding is required for efficient circularization. To further enhance P2 circularization, the 5′ exon was extended to include IBS2 in addition to IBS1. This modification substantially increased circularization efficiency, exceeding 50% at 60 min (Fig. 2D), and was therefore applied in subsequent constructs.

We explored various buffer compositions to optimize circularization efficiency. Specifically, we tested different combinations of monovalent (NH_4_Cl, NaCl or KCl) and divalent (MgCl_2_ or Mg(CH_3_COOH)_2_) salts, along with Tris pH 7.5 and DTT (Fig. 2E). Reactions were carried out at 55°C for 30 min. Notably, 1.5 M KCl combined with 50 mM Mg salt achieved higher circularization efficiency than either the modified SSB condition 1.5 M NH_4_Cl (Buffer A) or 1.5 M NaCl (Buffer I). Under the same conditions, MgCl_2_ showed slightly higher circularization efficiency than Mg(CH_3_COOH)_2_. Including both NH_4_Cl and KCl in the same reaction buffer did not enhance circularization; while maintaining the optimal concentration of KCl (1.5 M), reducing Mg from 50 mM to 10 mM completely abolished circularization (Fig. 2E). Taken together, Buffer K, composed of 1.5 M KCl, 50 mM MgCl_2_, 40 mM Tris-HCl (pH 7.5), and 5 mM DTT consistently yielded the highest circularization efficiency, achieving ∼85% circularization of CircLtrB-NLuc.

### NH_4_Cl- and KCl-based splicing conditions generate distinct RNA products

Early biochemical analyses showed that group II intron splicing remains active under high concentrations of monovalent salts and that salt identity strongly influences both splicing kinetics and products, with KCl leading to increased accumulation of free exons and reduced ligated exon formation compared to NH_4_Cl (Jarrell et al. 1988; Saldanha et al. 1999). More recent mechanistic work on group IIC introns further showed that NH_4_^+^ and K^+^ ions are not functionally interchangeable, with NH_4_Cl promoting intron conformations that support productive catalysis at elevated temperatures, whereas KCl biases reactions toward alternative pathways and altered intermediate profiles (Smathers and Robart 2020). Consistent with these prior reports, we found that NH_4_Cl and KCl-based splicing conditions also yield distinct outcomes in circLtrB PIE reactions. At both 30 and 60 min, KCl-based splicing conditions produced a higher circRNA-to-precursor ratio than NH_4_Cl, reaching ∼90% circularization, whereas NH_4_Cl-based reactions plateaued at ∼80% (Fig. 3A). However, by 60 min, reactions performed in KCl exhibited pronounced RNA smearing, suggesting RNA degradation (Fig. 3A). Accordingly, KCl-mediated circularization reactions were limited to 30 mins in subsequent experiments. These results suggest that, as in canonical group II intron splicing, salt identity in PIE reactions influences the balance between productive exon ligation and competing hydrolytic pathways, rather than simply affecting splicing efficiency.

**Figure 3.**
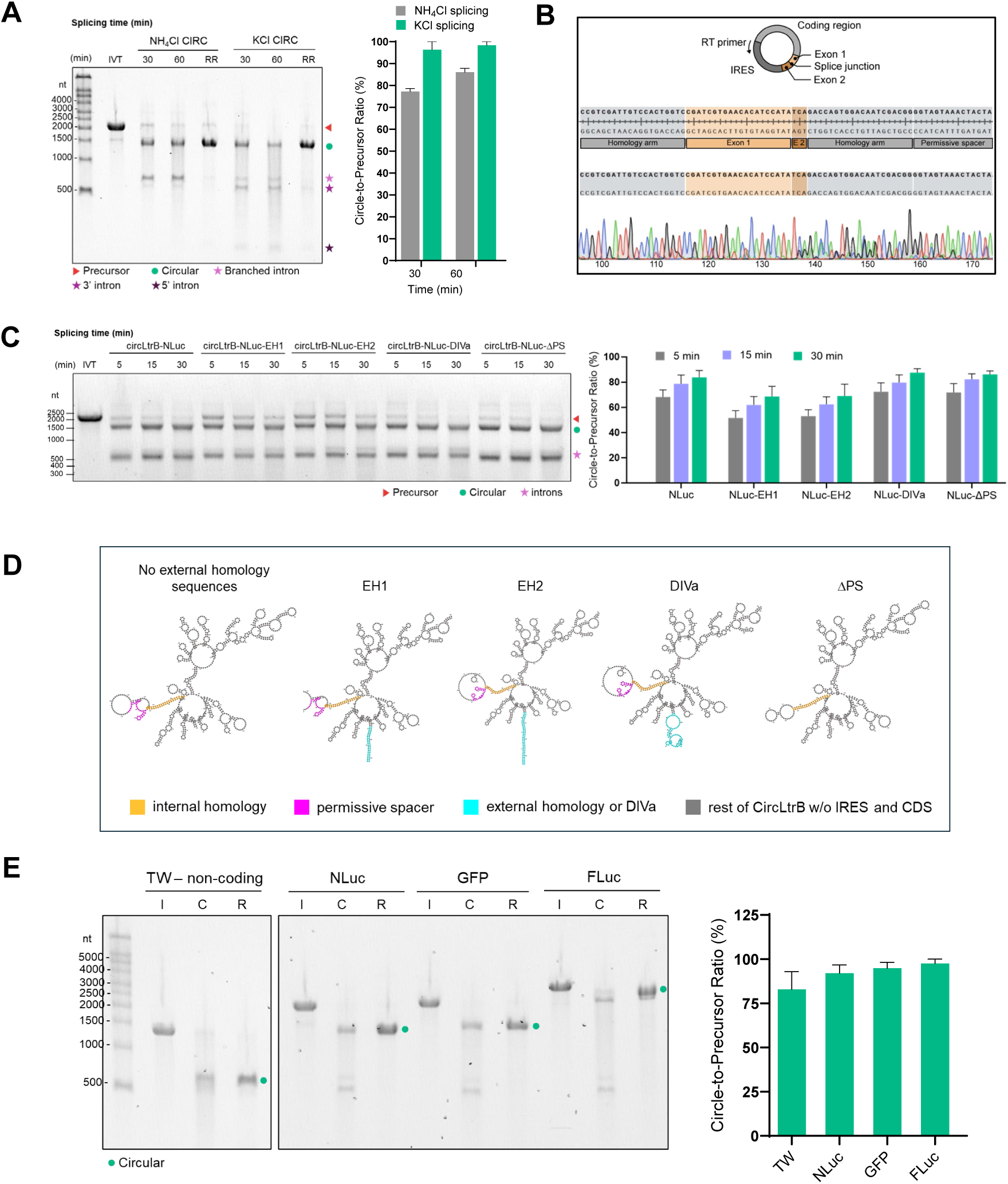
Design optimization of the PIE system and its capacity to circularize RNAs with varied payload sizes. *(A)* Circularization of CircLtrB-NLuc in NH_4_Cl- and KCl-based splicing buffers yields ∼80% and ∼90% circRNA, respectively (n = 3). Shown are the linear precursor (IVT), post-circularization product (CIRC), and RNase R–treated RNA (RR). *(B)* Representative Sanger trace from RT-PCR across the splice junction of RNase R–resistant RNA in *(A)*, confirming correct ligation. *(C)* Representative gel showing CircLtrB-NLuc splicing with cis-acting element variants predicted to enhance circularization (n = 3). *(D)* RNA secondary structure predictions for the constructs shown in *(C)* excluding the IRES and coding sequences were generated using RNAfold. *(E)* Representative gel showing self-splicing of CircLtrB constructs with different payload sizes (n = 3). Shown are the linear precursor (I), post-circularization product (C), and RNase R–treated RNA (R). Circularization percentages were calculated from purified circularization reactions (IVTC) and expressed relative to the remaining linear precursor.

To confirm that the product was indeed a covalently closed circRNA for both NH_4_Cl - and KCl-based circularization, the circularization products were subjected to RNase R digestion. RNase R efficiently degraded linear intermediates, while circRNAs resisted the RNase activity (Fig. 3A). Interestingly, we also observed RNase R-resistant species with size of ∼100 nt, potentially representing structured intermediates or byproducts. To validate the circular nature of the produced RNA, we performed RT-PCR using the RNase R-treated RNA as a template followed by Sanger sequencing across the backsplice junction. The resulting sequence aligned with the exon junction confirming the identity of the circRNA (Fig. 3B).

LtrB has previously been shown to undergo splicing facilitated by its intron-encoded maturase protein LtrA (Matsuura et al. 1997). We successfully expressed and purified the LtrA protein from *E. coli* using a resin batch purification method (Supplemental Fig. 2A). To confirm that it is functional, we first assayed its DNA endonuclease activity using a plasmid DNA containing the LtrB sequence as substrate. Following overnight incubation at 37°C, DNA cleavage was observed with 50 ng of LtrA, and complete digestion occurred with 100-300 ng, indicating that the purified LtrA is functionally active (Supplemental Fig. 2B). To assess maturase activity, we used the linear control Lin1 as a substrate under previously reported maturase-assisted circularization conditions, with incubations performed for 2 hr, 4 hr, or overnight at 30°C. As evidenced by the accumulation of the spliced exon band (Supplemental Fig. 2C), the purified LtrA protein is confirmed to have maturase activity. However, progressive smearing of RNA was observed at later time points, suggesting partial degradation. Next, we tested maturase-assisted circularization of CircLtrB-NLuc by incubating 100 nM circRNA precursor with increasing concentrations of purified LtrA (300, 600, or 900 nM) at 30°C for 1 hr to assess whether higher maturase levels would boost circularization efficiency. Simply, increasing LtrA concentration did not increase circularization efficiency. Densitometry analysis showed that the highest level of circRNA was observed at 300 nM LtrA, yielding ∼28% circular RNA relative to remaining linear precursor from 100 nM circLtrB-NLuc RNA (Supplemental Fig. 2D). To confirm circRNA species, these RNAs were purified and digested with RNase R. Linear RNA species were degraded, whereas the circular RNA remained intact (Supplemental Fig. 2E).

Notably, the linear RNA substrate used in these assays contains a truncated Domain 4 (D4), while D4 was split in the PIE construct. Because D4 encodes regions known to mediate critical interactions with the LtrA maturase (Matsuura et al. 2001), it is possible that the reduced splicing efficiency under maturase-assisted conditions may at least in part, reflect the absence of key D4 sequences or proper RNA folding required for optimal maturase binding and stabilization of the active intron conformation. Thus, self-splicing conditions were used in subsequent experiments.

### Outer homology arms do not enhance circularization efficiency in the circLtrB platform

To determine whether sequence context influences circularization efficiency in the circLtrB platform, we introduced different cis-acting sequence elements immediately upstream and downstream of the 5’ and 3’ splice junctions and evaluated their effects on PIE-mediated circularization. Across constructs, extending the splicing reaction time from 15 min to 30 min improved circRNA formation, indicating that circularization efficiency is partially time dependent under self-splicing conditions (Fig. 3C). Because group I intron-based circularization strategies enhanced efficiency by including external homology arms that promote exon alignment (Wesselhoeft et al. 2018), we tested two external homology designs (EH1 and EH2) that flanked the splice sites (Supplemental Table 1). Contrary to expectations, the external homology elements both reduced circular RNA formation (Fig. 3C-D), with circularization efficiencies decreasing to approximately ∼60% relative to constructs without the sequences (Fig. 3C). These findings show that, unlike group I intron platforms, circLtrB circularization is sensitive to local sequence context and does not benefit from exogenous homology elements, which may interfere with productive folding or exon alignment during group II intron-mediated splicing.

A cis-acting element that did not significantly reduce circularization efficiency was the DIVa interaction sites (Fig. 3C-D). In the LtrB group II intron, DIVa refers to a long-range tertiary interaction between domain IV (DIV) and domain I, which plays a key role in organizing the intron’s three-dimensional structure and positioning the splice sites during catalysis. DIVa contacts have been shown to stabilize intron folding and promote productive splicing by helping bring distal regions of the intron into close spatial proximity (Watanabe and Lambowitz 2004; Pyle 2016). Based on this mechanism, DIVa sequences were added to both ends of the PIE construct to test whether reinforcing DIV–DI interactions could reduce the effective distance between splice junctions and thereby enhance self-splicing efficiency (Supplemental Table 1, Fig. 3D). Consistent with this rationale, inclusion of DIVa sites did not measurably impair circularization (Fig. 3C), suggesting that DIVa-mediated tertiary interactions are compatible with the split intron architecture and do not interfere with productive PIE splicing.

Deletion of the permissive spacer also did not significantly reduce circularization efficiency (Supplemental Table 1, Fig. 3C-D). The deletion was explored as a strategy to reduce overall construct size and minimize extraneous sequences. However, the spacer was retained in subsequent designs, as it may influence RNA structure and translational performance, even in the absence of measurable effects on circularization.

### The circLtrB platform makes circRNAs with varying coding sequence lengths

To evaluate whether the size of the circRNA influences circularization efficiency in the circLtrB platform without external homology arms or DIVa, we compared circularization across constructs containing payloads of increasing size. Denaturing 1% E-Gel™ EX agarose gels (Fig. 3E) show efficient circRNA formation for all tested inserts, including a minimal non-coding sequence lacking any IRES (non-coding, 662 nt), nanoluciferase (NLuc, ∼1.3 kb), GFP (∼1.5 kb), and firefly luciferase (FLuc, ∼2.5 kb).

To evaluate the platform with larger constructs, we assessed the circularization of different Chikungunya antigen designs (∼3.9–4.6 kb) and an eGFP-tagged cystic fibrosis transmembrane conductance regulator (CFTR) transcript (∼6.1 kb) (Supplemental Fig. 3A-C, Supplemental Fig. 4A). These constructs were generated as part of separate projects. Under NH_4_Cl conditions, CircLtrB-CHIKV variants and CircLtrB-eGFP-CFTR were produced but with reduced efficiency (∼10%) and increased production of nicked RNA species (Supplemental Fig. 3A-C, Supplemental Fig. 4A). For comparison, the smaller CircLtrB-GFP construct was included and exhibited higher circularization efficiency, consistent with improved performance for shorter payloads (Supplemental Fig. 3A-B). Notably, the CHIKV circRNAs migrated above the precursor RNAs and their expected sizes, as indicated by RNase R-resistant bands, confirming their circular nature (Supplemental Fig. 3B). Because self-splicing conditions such as elevated salt concentrations and temperatures may promote degradation of longer RNAs, we also evaluated LtrA-assisted splicing. This approach yielded comparable levels of circLtrB-CHIKV but showed reduced release of intron intermediates, as indicated by less intense intron bands (Supplemental Fig. 3C). Given the increased sequence length and possible challenges in RNA folding and splice-site alignment, we revisited the addition of the external homology sequence (EH1); however, consistent with previous observations, this did not improve circularization efficiency.

Taken together, efficient circRNA formation was maintained across a wide range of coding sequence lengths, from minimal non-coding inserts to multi-kilobase protein-coding payloads, although efficiency decreased at longer transcript lengths.

### LtrB-derived circRNA support translation in mammalian cells

To assess how IRES selection influences translational output and persistence in this new circRNA platform, six viral IRES elements were evaluated using EGFP as a reporter. The IRESs tested included those derived from Aichivirus A, Crohivirus A, Coxsackievirus B3 (CVB3), Encephalomyocarditis virus (EMCV), Human Rhinovirus (HRV), and Salivirus A. These IRESs were incorporated into otherwise identical circLtrB constructs, enabling direct comparison of IRES-dependent translation from circLtrB. EGFP expression was monitored over a 4-day time course following transfection. All IRESs except Crohivirus A and Salivirus A expressed EGFP, with the CVB3 IRES showing the highest EGFP levels and therefore used in all subsequent expression studies (Fig. 4B). Consistent with our data, the CVB3 IRES has previously been shown to drive more potent cap-independent translation than other viral IRES elements in HEK293 cells, supporting its use for sustained protein expression in circRNA designs (Wesselhoeft et al. 2018).

**Figure 4.**
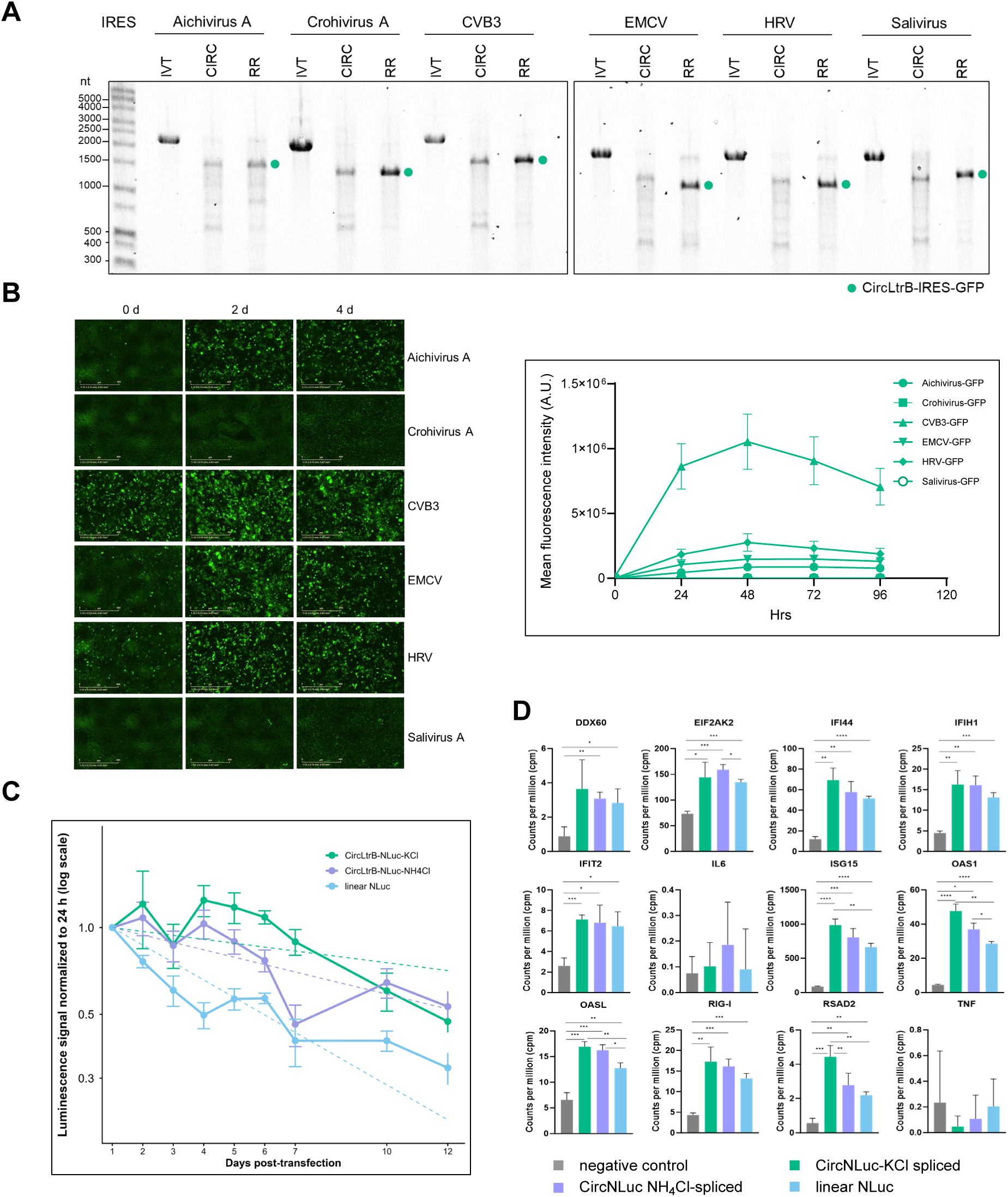
Translatability of CircLtrB constructs with different IRESs and comparison of linear RNA versus circLtrB translation and immune marker activation. *(A)* Representative gel showing self-splicing of CircLtrB-GFP constructs with different IRES. Shown are the linear precursor (IVT), post-circularization product (CIRC), and RNase R–treated RNA (RR). *(B)* HEK293 cells were transfected with CircLtrB-GFP circRNA constructs with the indicated IRES. GFP fluorescence was monitored over a 4-day time course, and mean fluorescence intensity (MFI) was quantified at each time point (n = 6 technical replicates, 5 fields per replicate) over the 4-day time course (n = 6). Scale bar indicated on image. *(C)* Luminescence signal from HEK293 cells transfected with CircLtrB-NLuc under KCl (green) or NH_4_Cl (purple) conditions, or a linear NLuc RNA (light blue), measured over 12 days post-transfection. Signals were normalized to the 24 h time point for each condition and plotted on a log scale. Dashed lines represent fitted decay curves. Data are shown as mean ± SEM (n = 3). *(D)* RNA expression levels of selected immune marker genes in U2OS cells transfected with negative control (linear mCherry RNA only), linear NLuc, or circLtrB constructs generated under NH_4_Cl or KCl splicing conditions and co-transfected with linear mCherry RNA.

To compare expression from linear NLuc mRNA and circLtrB-NLuc generated under KCl or NH_4_Cl conditions, we transfected HEK293 cells and monitored protein levels over a 12-day period. Linear NLuc expression peaked at 24 h post-transfection and declined over time while circRNA-derived NLuc expression was more sustained (Supplementary Fig. 5). Although absolute NLuc levels from linear mRNA remained higher at later time points, the signal declined more rapidly compared to circRNA. By comparison, circLtrB-NLuc exhibited lower initial expression but also yielded more stable signals over extended times, resulting in a more persistent expression profile (Fig. 4C, Supplementary Fig. 5). Minor fluctuations in signal over time are likely attributable to experimental factors such as cell perturbation during media changes and sampling. Also, the comparatively lower expression from circRNA likely reflects sequence-dependent factors affecting translatability, such as optimized spacer elements or local sequence context surrounding the IRES, which were not comprehensively explored in this study.

Given the complexity of expressing the transmembrane protein CFTR, we were particularly interested in whether circLtrB could faithfully express such a challenging protein (Lukacs and Verkman 2012). We transfected purified and phosphatase-treated CircLtrB-eGFP-CFTR into Calu-3 lung epithelial cells and monitored GFP expression. eGFP-CFTR localized to the cell membrane at 24 hr post-transfection and remained detectable at 70 hr, demonstrating that the circRNA is functional despite lower circularization efficiency (Supplemental Fig. 4A-B).

### RNA-seq reveals similar innate immune responses to linear and circLtrB-derived circRNAs

To compare innate immune responses elicited by circRNA versus linear capped mRNA, RNA-seq was performed on cells transfected with circLtrB-NLuc generated under either NH_4_Cl or KCl splicing conditions, alongside a linear NLuc-3X-FLAG mRNA control. All RNA preparations were phosphatase-treated prior to transfection to minimize contributions from residual linear precursor or RNase R digestion byproducts. Samples were co-transfected with linear mCherry mRNA to control for transfection efficiency, and mCherry-only transfections served as a negative control. Library construction, sequencing, quality control, and read count quantification were performed by Plasmidsaurus. Samples were strongly correlated, with most uniquely mapped reads corresponding to protein-coding genes, indicating high data quality and consistency (Supplemental Fig. 6A-B).

Differential expression analysis revealed that both circLtrB-NLuc and linear NLuc mRNA induced upregulation of innate immune-associated genes relative to the mCherry only control (Supplemental Fig. 6C). Induced transcripts included canonical interferon-stimulated genes and RNA-sensing pathway components, such as DDX60, EIF2AK2 (PKR), IFI44, IFIH1 (MDA5), IFIT2, IL6, ISG15, OAS1, OASL, RIG-I, RSAD2, and TNF, with ISG15 exhibiting the highest level of induction across conditions (Fig. 4D).

Notably, the overall immune activation profile elicited by circLtrB-NLuc was comparable to that induced by linear NLuc mRNA, with some markers including IFIH1, ISG15, OAS1, OASL and RSAD2 showing statistically significant higher expression in circLtrB-transfected cells. This observation is consistent with prior work demonstrating that exogenous circRNAs generated *in vitro* can potently stimulate innate immune gene expression and that mammalian cells distinguish self from non-self circRNAs based on intron identity and sequence origin (Chen et al. 2017; Wesselhoeft et al. 2019; Liu et al. 2022; Jiang et al. 2024). In this context, the circLtrB constructs contain group II intron-derived sequences of bacterial origin in addition to the permissive spacer and polyA-C sequences, which likely represent non-self RNA features that may contribute to immune recognition. On the other hand, the linear NLuc-3X-FLAG mRNA contains engineered 5’ UTR and 3’ UTR sequences, and a 3’ poly(A) tail. While these elements may still elicit immune responses, they are likely less immunostimulatory than bacterial-derived intron sequences (Liu et al. 2022; Tong et al. 2025b).

Moreover, a possible explanation for the comparable immune signatures between circRNA and linear RNA is that activation may be driven in part by RNA byproducts such as double-stranded RNA (dsRNA) formed during transcription, which can engage cytosolic RNA sensors (Kato et al. 2006). In addition, with the exception of RSAD2, no other significant differences in immune gene induction were observed between NH_4_Cl or KCl-spliced circLtrB-NLuc, indicating that splicing buffer composition does not greatly influence innate immune activation at the transcriptomic level (Fig. 4D). IL6 and TNF levels also remained low across all samples, suggesting that while there is activation of antiviral and interferon-stimulated gene pathways, there is no induction of broad pro-inflammatory cytokine responses.

Together, these results indicate that circLtrB-derived circRNAs engage host innate immune pathways like linear mRNA under the conditions tested, and that this activation is driven by RNA byproducts such as dsRNA rather than splicing conditions.

## MATERIALS AND METHODS

### Plasmid construction and *in vitro* transcription templates

Target sequences were codon-optimized for *E.coli* or mammalian expression, as appropriate, using the GeneArt Gene Synthesis tool (Thermo Fisher Scientific). Optimized sequences were synthesized and cloned into GeneArt’s high-expression vectors and delivered as complete, sequence-verified plasmids. Site directed mutagenesis using the QuikChange Lightning Site-Directed Mutagenesis Kit (Agilent, Cat.: 210519) was performed to eliminate the early T7 pause recognition site while retaining the loop structure using primers (Supplemental Table 1). CircLtrB-GFP constructs with distinct IRES elements were generated by Golden Gate assembly using BsaI-HFv2 (NEB, Cat.: R3733S) and T4 DNA ligase (NEB, Cat.: M0202S). Constructs with alternative cis-acting elements and coding sequences were assembled using the KLD Enzyme Mix (NEB, Cat.: M0554S) or NEBuilder HiFi DNA Assembly (NEB, Cat.: E2621S), as appropriate. Ligation products were transformed into *E. coli* DH5α chemically competent cells (NEB, Cat.: C2987U) and plated on LB agar containing ampicillin. Correct plasmids were verified by full plasmid sequencing (Plasmidsaurus), followed by large-scale plasmid DNA preparation using the ZymoPURE II Plasmid Maxiprep Kit (Zymo Research, Cat.: D4203). To prepare *in vitro* transcription templates, purified plasmids were subsequently linearized with BspQI restriction enzyme (NEB, Cat.: R0712S), and the resulting digests were purified using the DNA Clean & Concentrator-25 kit (Zymo Research, Cat.: D4033) according to the manufacturer’s instructions. Plasmids used in this manuscript are available on Addgene under accession numbers ##########.

### LtrA expression and purification

The LtrA open reading frame was codon-optimized (GeneArt, Thermo Fisher Scientific) and cloned into an *E. coli* expression vector with N-terminal 6xHis and V5 epitope tags. The resulting plasmid was transformed into BL21 Star (DE3) cells (NEB, Cat.: C2527H). Starter cultures (50 mL) were grown overnight in LB supplemented with 50 μg/mL ampicillin and used to inoculate 1 L Terrific Broth (TB). Cultures were grown at 37°C with shaking at 200 rpm until reaching an OD_600_ of 0.3-0.4, followed by incubation at 18°C for 1 h prior to induction. Protein expression was induced by addition of IPTG to a final concentration of 1 mM, and cultures were incubated overnight at 18°C. Cells were harvested by centrifugation for 10 min at 5,000 x g using a Beckman Coulter J-LITE JLA-8.1000 fixed-angle rotor and resuspended in 30 mL pre-chilled lysis buffer (50 mM Tris-HCl, 1 M NaCl, 30 mM imidazole, 10% (v/v) glycerol, pH 8.0). Cell disruption was performed by sonication using a Pulse 150 UltraSonic Homogenizer (Benchmark Scientific, Cat.: DP0150) equipped with a 6 mm horn, applying two cycles with 35% amplitude, 5 s ON / 5 s OFF, for a total sonication time of 3 min, while maintaining samples on ice.

Lysates were clarified by centrifugation at 30,000 x g for 30-45 min at 4°C using a Beckman Coulter JA-25.50 fixed-angle rotor. The cleared lysate was loaded onto a 5 mL HisTrap HP column (Cytiva, Cat.: 17-5248-01) using an ÄKTA Start system, followed by washing with 80 mL lysis buffer. Bound proteins were eluted by gradient elution (0–100%) over 50 mL elution buffer (50 mM Tris-HCl, 500 mM NaCl, 500 mM imidazole, 10% (v/v) glycerol, pH 8.0). Eluted fractions containing LtrA were pooled and dialyzed overnight at 4°C using Spectra/Por 6 regenerated cellulose dialysis tubing (MWCO 3.5 kDa; Spectrum) against 1 L dialysis buffer (50 mM Tris-HCl, 500 mM NaCl, 10% (v/v) glycerol, pH 8.0). Protein concentration was determined using a DeNovix spectrophotometer. Protein expression and purity were assessed by SDS-PAGE and western blotting using an anti-V5 antibody (Invitrogen, Cat.: MA5-15253). Purified LtrA protein was aliquoted, flash-frozen, and stored at -80°C until use.

### Self and maturase assisted splicing of LtrB linear intron and PIE variants

RNA was generated by *in vitro* run-off transcription using the HiScribe T7 High Yield RNA Synthesis Kit (NEB, Cat.: E2040S) and BspQI-HF (NEB, Cat.: R3712S) linearized DNA plasmid as template. Reactions were incubated for 2 h and 30 min at 37°C with shaking at 250 rpm, and the RNA product was column purified using the Monarch RNA Cleanup Kit (NEB, Cat.: T2050S). To initiate self-splicing, RNA was incubated at 70°C for 3 min, then incubated in self-splicing buffer (SSB, 0.5/1.5 M NH_4_Cl, 0.1/1.5 M KCl, 10/20/50 mM MgCl_2_ or Mg(CH_3_COOH)_2_, 40/50 mM Tris–Cl, pH 7.5, 5 mM DTT) at 37°C or 55°C. The best-optimized reaction condition (SSB: 1.5 M KCl, 50 mM MgCl_2_, 40 mM Tris–Cl, pH 7.5, 5 mM DTT at 37°C for 15 min) was selected for further experiments. For maturase splicing, 100 nM IVT RNA was incubated in maturase-assisted circularization buffer (MACB, 500 mM NH_4_Cl, 10 mM MgCl_2_, 40 mM Tris pH 7.5, 5 mM DTT, 0.5 U/µl SUPERase·In™ RNase Inhibitor) with varying amounts of purified LtrA (300 nM, 600 nM, or 900 nM) at 30°C. Circularization reactions were optimized to perform best at final RNA concentrations of 0.1–1 µg/µL without agitation. The circularization reaction was column purified using a Monarch RNA Cleanup Kit.

### Verification of circRNA

To remove linear RNA contaminants and to confirm that the resulting splicing product is a closed circle, the purified circularization products were treated with 1 U of RNase R (LGC Biosearch Technologies, Cat.: RNR07250) per 1 μg of circularized RNA at 37°C for 1 h with 500 rpm shaking. Reactions were stopped at indicated times and were column-purified using the RNA Clean & Concentrator 5 µg (Zymo Research, Cat.: R1014) Furthermore, the band corresponding to the circRNA was collected by running into a 1% E-Gel™ EX Double Comb Agarose Gel (Thermo Fisher Scientific, Cat.: A42345) until the desired band reached the second-tier wells. This was used as a template for the cDNA synthesis using the Maxima H Minus First Strand cDNA Synthesis Kit (Thermo Fisher Scientific, Cat.: K1681) with a CVB3-gene-specific reverse primer (Supplement Table 2). Three microliters of the cDNA synthesis reaction were used as a PCR template to amplify the region containing the backsplice junction with primers flanking the junction (Supplement Table 2). The PCR product was column purified (Zymo Research, Cat.: D4014) and was sequenced by the Sanger method (Eurofins).

To assess RNA integrity, samples were denatured prior to electrophoretic analysis. For visualization on a 1% E-Gel™ EX agarose gel (Thermo Fisher Scientific, Cat.: G401001), RNA was denatured in 50% (v/v) formamide at 65°C for 3 min. For analysis on a 1.2% FlashGel® cassette (Lonza, Cat.: 57027), RNA was denatured in 50% (v/v) formaldehyde buffer (Lonza, Cat. 50571) under identical conditions. Following denaturation, samples were immediately cooled on ice for 2 min and loaded alongside the Millennium™ RNA Marker (Invitrogen, Cat.: AM7150).

### Live fluorescence microscopy imaging

HEK293 cells were cultured in Dulbecco’s Modified Eagle Medium (Gibco, Cat.: 11965092) supplemented with 10% Tet-free FBS (Thermo Fisher Scientific, Cat.: A4736301) (DMEM + 10% Tet-free FBS) at 37°C and seeded into a 96-well plate (Corning, Cat.: 3596) at a target density of 4 × 10^4^ cells per well. RNA was transfected at a total amount of 150 ng (140 ng CircLtrB-IRES-GFP RNA + 10 ng linear mCherry RNA) per well using the Lipofectamine™ MessengerMAX™ Transfection Reagent (Thermo Fisher Scientific, Cat.: LMRNA015) according to the manufacturer’s instructions. Each condition was performed in six independent replicate wells. Live-cell fluorescence imaging using the Incucyte S3 Live-Cell Analysis System (Sartorius) was performed to monitor and quantify CircLtrB-GFP expression over time. For each well, five non-overlapping fields of view were acquired at the indicated time points. The signals for each image were quantified using the Incucyte S3 software (Incucyte 2022B Rev2), averaged for each well, and are shown as the mean fluorescence intensity.

### Calu-3 cell culture and fluorescence imaging

Human lung epithelial Calu-3 cells (ATCC, Cat.: HTB-55) were maintained in Eagle’s Minimum Essential Medium (EMEM; ATCC, Cat.: 30-2003) supplemented with 10% FBS and maintained in a 37°C incubator with 5% CO_2_. The circLtrB-eGFP-CFTR plasmid template was synthesized and provided by a commercial vendor (EZYVEC, Sartorius Polyplus). Purified circLtrB-eGFP-CFTR RNA were phosphatase-treated with Quick CIP (NEB, Cat.: M0525S) prior to transfection. Cells were co-transfected with circLtrB-eGFP-CFTR RNA and mCherry mRNA per well using the Lipofectamine™ MessengerMAX™ Transfection Reagent according to the manufacturer’s instructions. Media were replaced 4 h post-transfection, and cells were imaged 24 h post-transfection using the Echo Revolve Fluorescence Microscope (Echo Laboratories) at 10X magnification. GFP and mCherry signals were captured using appropriate filter sets, and images were pseudo-colored (GFP, blue; mCherry, yellow) for visualization.

### Nanoluciferase reporter assay

HEK293 cells were cultured in DMEM + 10% Tet-free FBS at 37°C and seeded in white 96-well plates (Perkin Elmer, Cat.: 101677-210) at a target density of 4 × 10^4^ cells per well. RNA was transfected at a total amount of 150 ng (140 ng CircLtrB-NLuc or linear NLuc RNA + 10 ng linear mCherry RNA) per well using the Lipofectamine™ MessengerMAX™ Transfection Reagent according to the manufacturer’s instructions. Each condition was performed in three independent replicate wells. Media were replaced 4h after transfection and refreshed daily thereafter. Samples were assayed at 8 h and at 1, 2, 3, 4, 5, 6, 7, 10, 12 days post-transfection for NanoLuc luciferase activity using the Nano-Glo® Dual-Luciferase Reporter Assay System (Promega, Cat.: E2920), with luminescence quantified using a SpectraMax iD5 microplate reader (Molecular Devices).

### RNA extraction and bulk RNA sequencing

U2OS cells were cultured in in McCoy’s 5A plus 10% FBS and maintained in a 37°C incubator with 5% CO_2_. Linear NLuc mRNA and circLtrB-NLuc RNAs were phosphatase-treated with Quick CIP prior to transfection and delivered at equimolar concentrations, using Lipofectamine™ MessengerMAX™ Transfection Reagent according to the manufacturer’s instructions, together with a linear mCherry mRNA to serve as transfection control. Cells transfected with linear mCherry mRNA served as the negative control. The transfection medium was removed, and fresh complete medium was added to the cells 4 h after transfection.

Total RNA was extracted 24 h post-transfection using the Monarch Spin RNA Isolation Kit (NEB, Cat. No. T2110S) according to the manufacturer’s instructions. RNA quality and integrity were assessed using an Agilent 4200 TapeStation (Agilent Technologies). Purified RNA samples were submitted to Plasmidsaurus, Inc. (Eugene, OR) for bulk RNA sequencing. Library preparation and sequencing were performed by Plasmidsaurus according to their standard protocols. Raw FASTQ files and vendor-generated preliminary analysis reports and figures were returned. The RNA-sequencing data generated by Plasmidsaurus in this study have been deposited in the NCBI Gene Expression Omnibus (GEO) under accession number GSE330383. Raw sequencing reads were assessed for quality using FastQC v0.12.1. Reads were then processed with fastp v0.24.0, which included poly-X tail trimming, 3′ quality-based trimming, filtering for a minimum Phred quality score of 15, and removal of reads shorter than 50 bp. Quality-filtered reads were aligned to the reference genome using STAR aligner v2.7.11, with non-canonical splice junctions removed and unmapped reads retained. Aligned reads were coordinate-sorted using samtools v1.22.1. PCR and optical duplicates were removed via UMI-based deduplication using UMIcollapse v1.1.0. Alignment quality metrics, strand specificity, and read distribution across genomic features were evaluated using RSeQC v5.0.4 and Qualimap v2.3. All quality control metrics were aggregated into a comprehensive report using MultiQC v1.32. Gene-level expression quantification was performed using featureCounts (subread package v2.1.1) with strand-specific counting, fractional assignment of multi-mapping reads, and exons and 3′ untranslated regions (UTRs) specified as features, grouped by *gene_id*. Gene counts were annotated with gene biotype and additional metadata extracted from the reference GTF file. Normalized expression values were calculated using trimmed mean of M-values (TMM) normalization. Sample-sample correlations were computed using Pearson correlation and visualized via sample correlation heatmaps and principal component analysis (PCA). Differential expression analysis was performed using edgeR v4.0.16, following standard workflows including filtering of lowly expressed genes with edgeR::filterByExpr using default parameters. Where applicable, functional enrichment analysis was conducted using gene set enrichment analysis (GSEA) with gseapy v0.12, using the MSigDB Hallmark gene sets.

## AUTHOR CONTRIBUTIONS

Kathrina D. Castillo: Conceptualization, Formal analysis, Methodology, Validation, Investigation, Data curation, Visualization, Supervision, Writing-original draft. Chotiwat Seephetdee: Methodology, review & editing. Wataru Horikawa: Formal analysis, Methodology. Daniel Kiss: Conceptualization, Methodology, Formal Analysis, Writing – original draft, Writing - review & editing, Resources, Funding Acquisition, Supervision.

## ACKNOWLEDGEMENTS

We would like to thank all the other members of the Kiss Lab and the Houston Methodist Research Institute (HMRI) Center for RNA Therapeutics for their support and fruitful discussions regarding the project. We would like to thank the staff of the Advanced Cellular and Tissue Microscopy Core Facility and the RNACore at the HMRI for technical support. We thank Rachel Whitehead from HMRI for assistance with figure illustrations and organization for select figures in Figures 1-3.

The content presented here is solely the responsibility of the authors and does not represent the official views of the HMRI or CEPI.

## DATA DEPOSITION

RNA-sequencing raw FastQ files and processed text files were deposited at GEO with accession numbers GSE330383 and are publicly available as of the date of publication.

## SUPPLEMENTARY FIGURE LEGENDS

**Supplement Figure 1.**
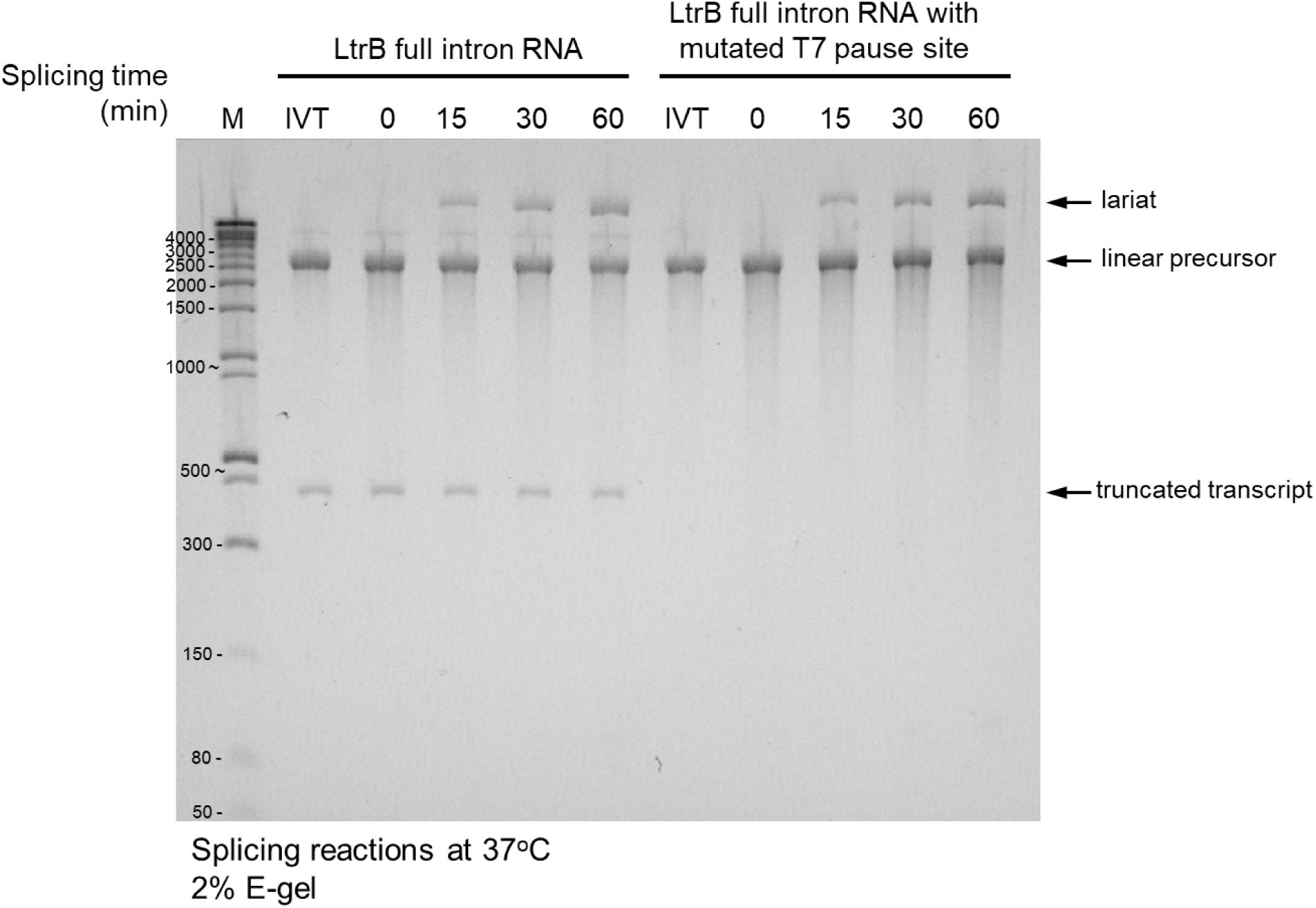
Removal of the T7 early termination site in LtrB Domain 1 (D1). Self-splicing of the 2.5-kb linear LtrB intron RNA was assessed at the indicated time points and resolved on a 2% E-Gel™ agarose gel. Reactions were performed under standard conditions (0.5 M NH_4_Cl, 50 mM MgCl_2_, 40 mM Tris-HCl pH 7.5, 5 mM DTT, 1 U/μL SUPERase•In; 37°C). A ∼400-nt truncated species observed with the native LtrB sequence is eliminated upon mutation of the T7 early termination site in D1.

**Supplement Figure 2.**
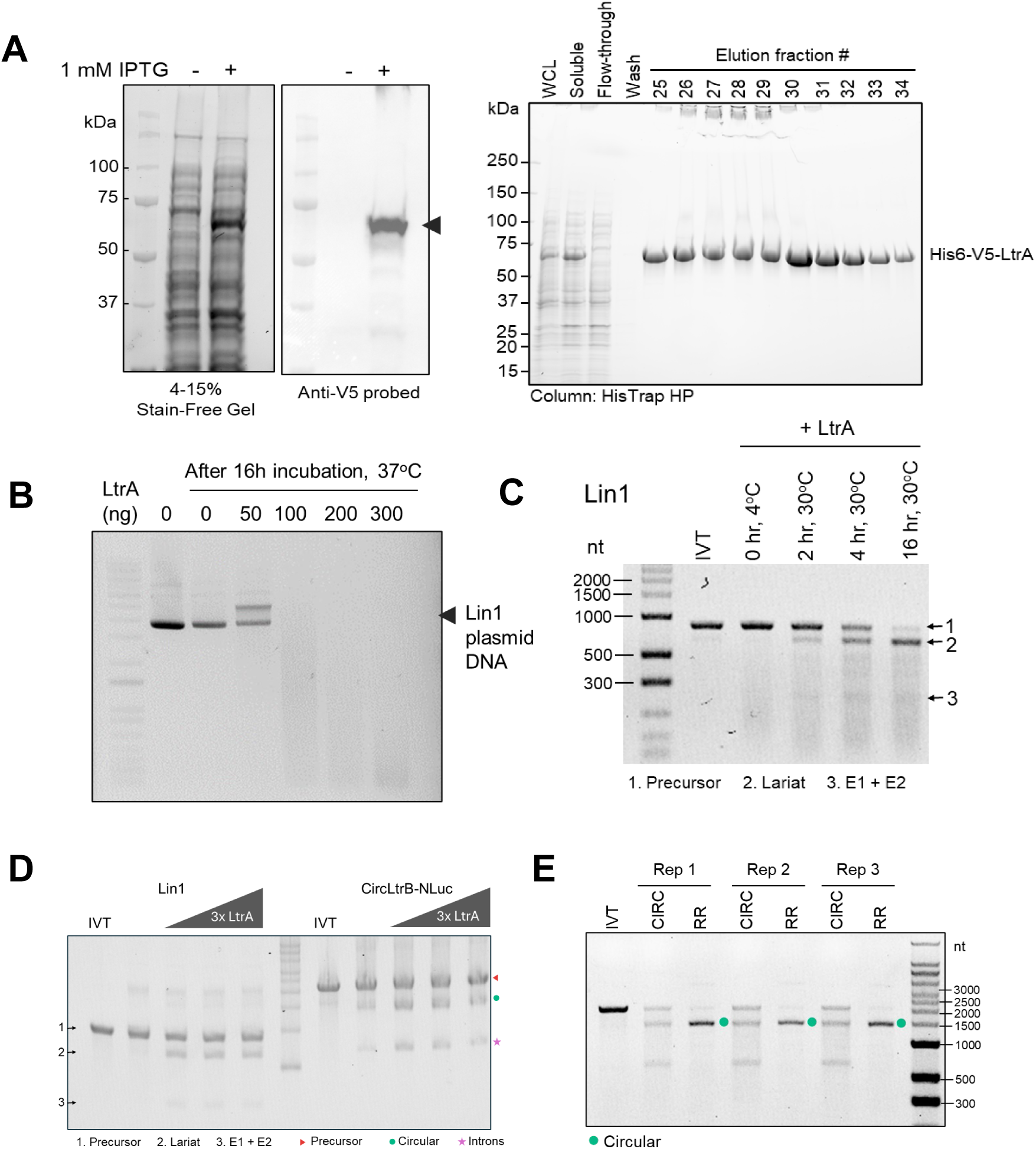
Assessment of LtrA maturase-assisted splicing in the circLtrB system. *(A)* Purification of 6x His-V5–LtrA. The expected ∼73.9-kDa band is indicated by the black arrow. *(B)* Endonuclease activity of purified LtrA assessed using a LtrB-containing plasmid DNA substrate. *(C)* Splicing time course for linear LtrB intron Lin1 showing the precursor (826 nt) and spliced intron/lariat (638 nt)*. (D)* LtrA-dependent splicing of the linear intron (Lin1) and CircLtrB-NLuc at increasing LtrA concentrations (0, 300, 600, 900 nM) with 100 nM RNA. *(E)* RNase R–resistant signal confirms circular RNA formation from LtrA-mediated splicing of LtrB-NLuc (green dot). Reactions were performed in maturase-assisted circularization buffer (MACB; 0.5 M NH_4_Cl, 10 mM Mg^2^⁺) at 30°C for 1 hr.

**Supplement Figure 3.**
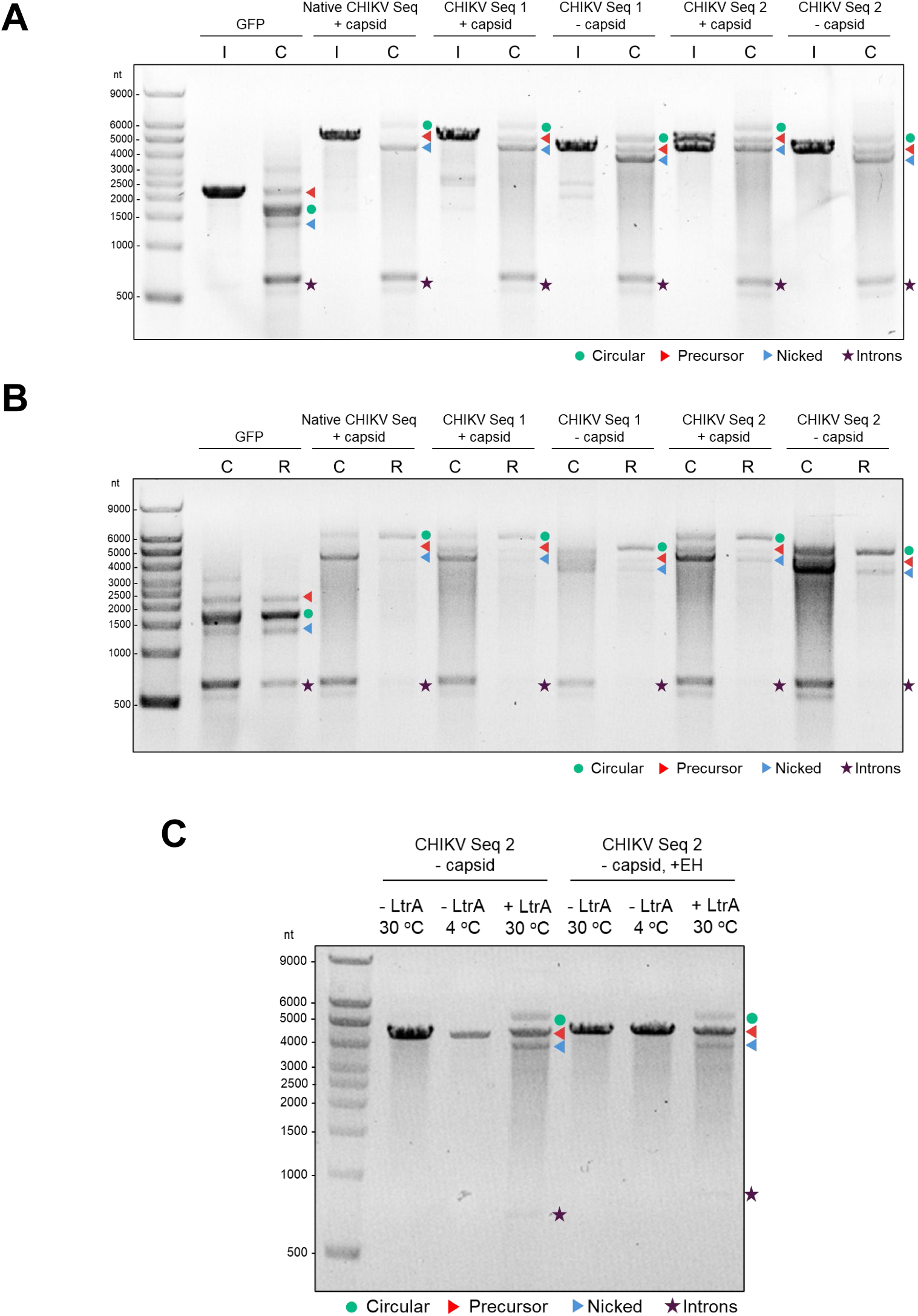
circLtrB-derived circRNA support circularization of Chikungunya virus (CHIKV) antigen variants. *(A)* Circularization of circLtrB-GFP and CHIKV antigen variants (native with capsid, and sequence designs 1 and 2, with or without capsid) under self-splicing conditions. Shown are the linear precursor (I) and post-circularization (C) RNA. *(B)* Post-circularization (C) and RNase-R treated RNA (R) of precursor RNAs shown in (*A)*. *(C)* LtrA-dependent splicing of sequence design 2 without capsid precursor RNA with or without external homology sequences. Reactions were performed in maturase-assisted circularization buffer (MACB; 0.5 M NH_4_Cl, 10 mM Mg²⁺) at 30°C for 15 min. Reactions without LtrA were incubated at 30°C or 4°C for 15 min to control for buffer-dependent splicing.

**Supplement Figure 4.**
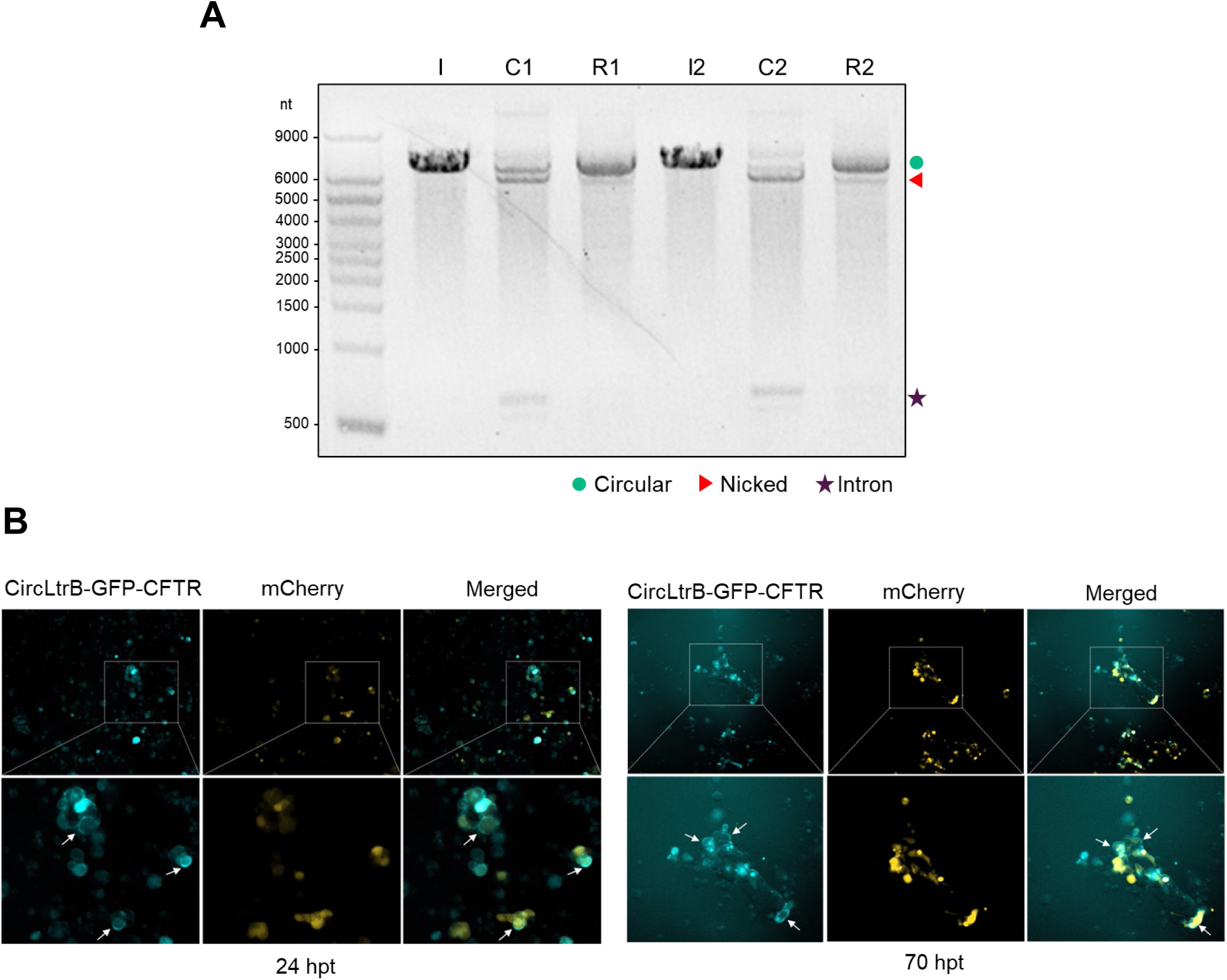
circLtrB-derived circRNA support functional protein expression, including membrane-localized proteins. *(A)* Circularization of circLtrB-eGFP-CFTR from two independent reactions (n = 2). Shown are the linear precursor (I), post-circularization product (C), and RNase R–treated RNA (R). *(B)* circLtrB-eGFP-CFTR was co-transfected into Calu-3 cells with mCherry, showing membrane-localized GFP expression at 24 h and 70 h post-transfection. Images were acquired on an ECHO microscope (10X). GFP and mCherry are pseudo-colored blue and yellow, respectively.

**Supplement Figure 5.**
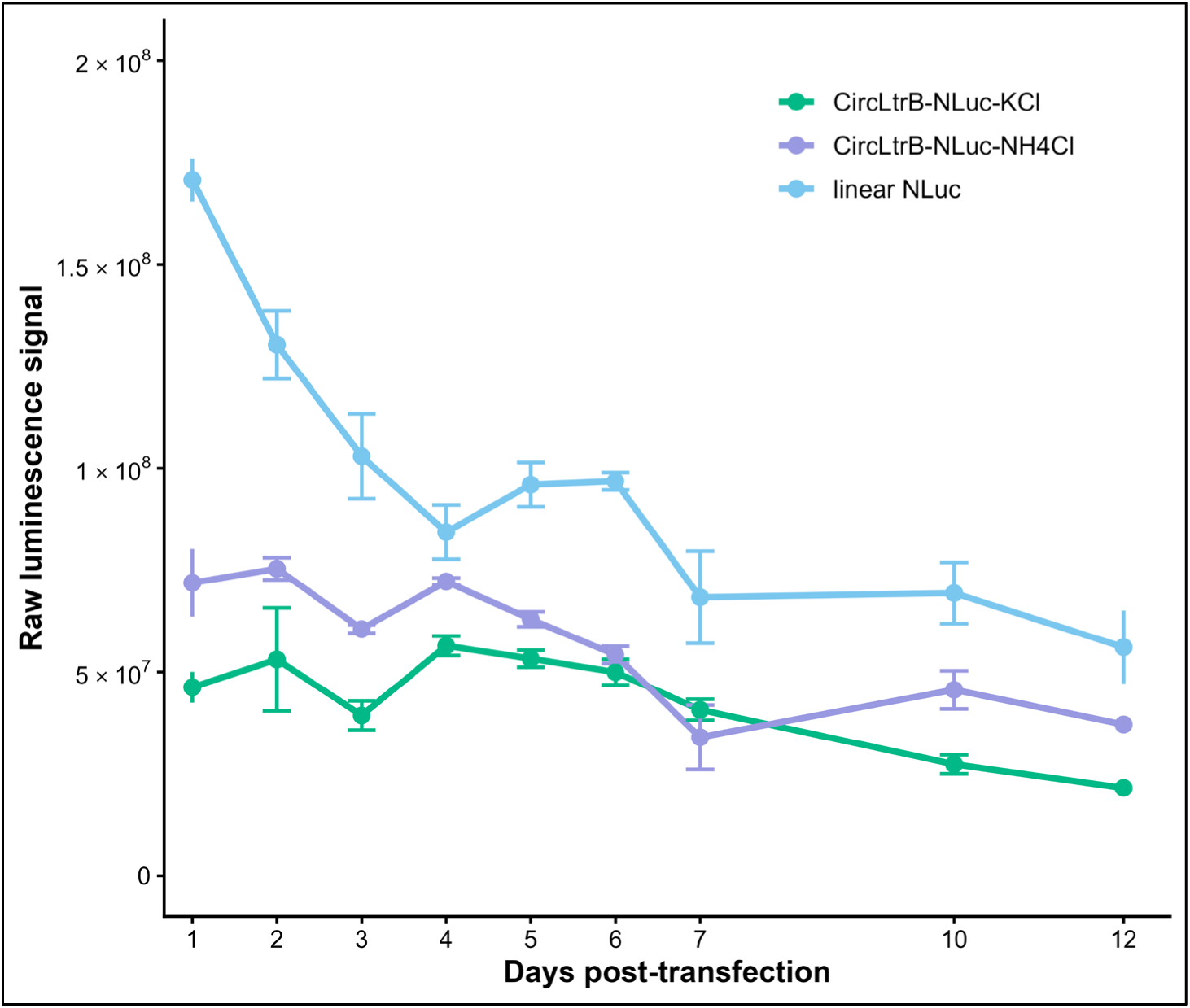
Time-course of NLuc expression from linear and circLtrB-derived RNAs. Luminescence signal from HEK293 cells transfected with CircLtrB-NLuc under KCl (green) or NH_4_Cl (purple) conditions, or a linear NLuc RNA (light blue), measured over 12 days post-transfection. Data are shown as raw luminescence values (mean ± SEM, n = 3).

**Supplement Figure 6.**
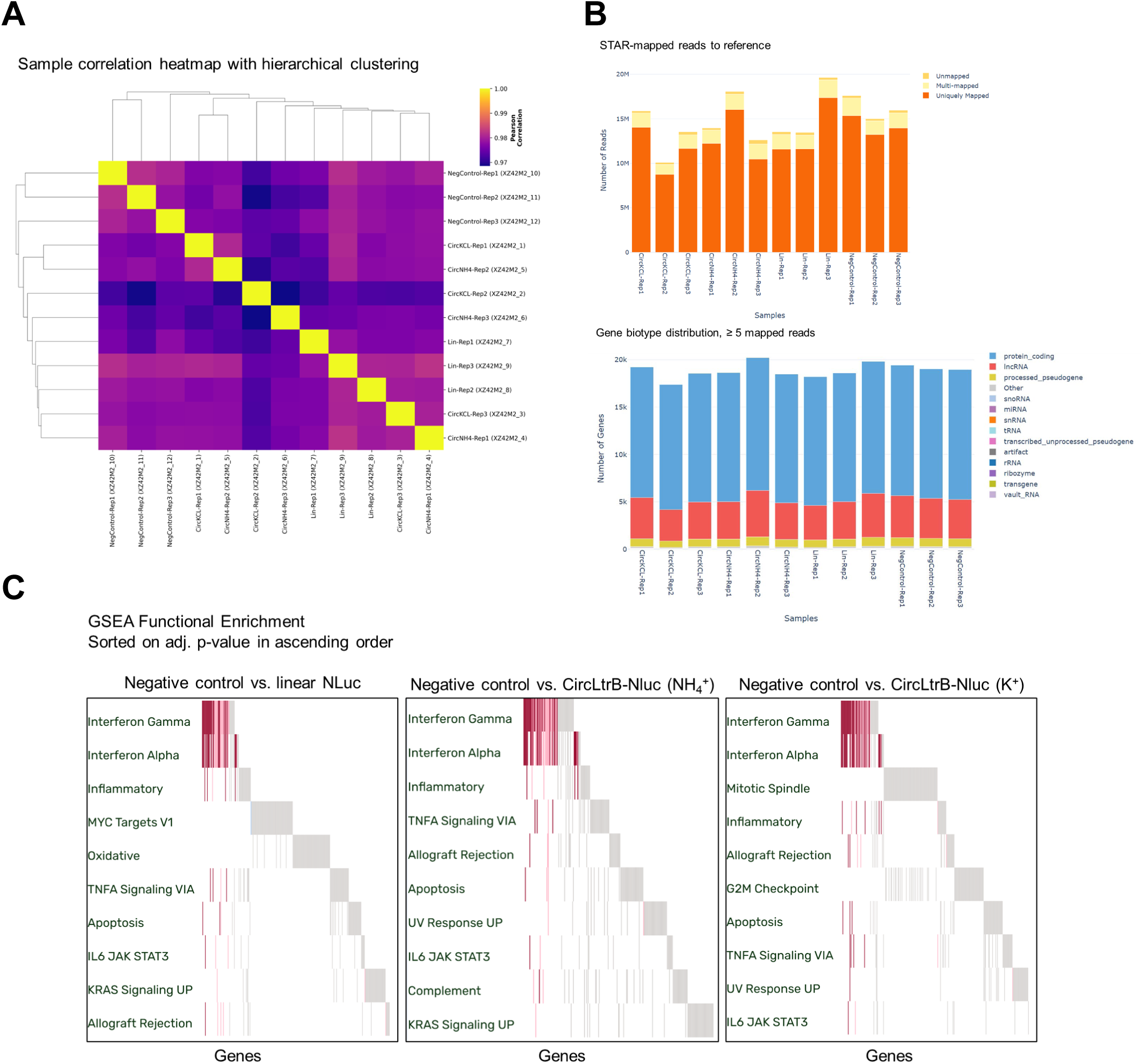
RNA-sequencing quality assessment and functional enrichment analysis. *(A)* Sample correlation heatmap showing pairwise correlations between RNA-seq samples with hierarchical clustering based on global gene expression profiles. *(B)* Summary of sequencing read alignment, showing the number of reads mapped to the reference genome using the STAR aligner (top) and the distribution of uniquely mapped reads by gene type (bottom). *(C)* Gene Set Enrichment Analysis (GSEA) of differentially expressed genes comparing the negative control condition (cells transfected with linear mCherry only) versus test RNA conditions (cells transfected with either linear NLuc or CircLtrB-Nluc RNA, each co-transfected with linear mCherry RNA as a transfection control). Enriched gene sets are sorted by adjusted p-value (false discovery rate, FDR) in ascending order.

